# Characterization of the human fetal rete region by single cell transcriptional analysis of gonads and mesonephros/epididymis

**DOI:** 10.1101/2022.07.20.500903

**Authors:** Jasin Taelman, Sylwia M. Czukiewska, Ioannis Moustakas, Yolanda W. Chang, Sanne Hillenius, Talia van der Helm, Hailiang Mei, Xueying Fan, Susana M. Chuva de Sousa Lopes

## Abstract

During development of the male reproductive tract, the human rete functions as a bridging structure between the seminiferous tubes and the efferent ducts of the epididymis. However, despite its significant contribution to human sex-specific gonadogenesis and future male fertility, the rete testis remains poorly understood. To investigate fetal rete testis development, we have performed single-cell transcriptomics on human fetal testes (and ovaries) and mesonephros/epididymis from first and second trimester. By revealing KRT19 and PAX8 as rete markers, we were able to identify the molecular signatures of the rete epithelial cells and mesonephros/epididymis epithelial cells. In the process, we have identified a population of Sertoli cells and germ cells in the rete testis. Moreover, we also revealed a small population of epithelial cells in the rete ovarii, comparable to the epithelial cells in the rete testis. Together, our study provides insights into human fetal sex-specific gonadogenesis and development of the male reproductive tract beyond the gonads.

## INTRODUCTION

Understanding human gonadogenesis and the role of the associated somatic niche remains limited due to tissue inaccessibility and ethical restrictions. Although certain aspects of human germ cell development and the contribution of the gonadal somatic niche are now relatively well characterized, even in term of transcriptional profile, limited attention has been given to the rete region. In both male and female gonads, both the cellular identity and sex-specific progression remains obscure. However, the rete region is integral to gonadal development and function in both sexes. In adult testes, the rete testis forms a crucial connection between the seminiferous tubes, containing the gonadal germ cell niche, and the extragonadal efferent ducts of the epididymis, developed from mesonephric tubules early during development (Combes et al., 2009, Major et al., 2021, Malolina and Kulibin, 2017, Malolina and Kulibin, 2019). In contrast to males, the primary female rete network regresses over time, yet cells from the ovarian rete have been associated with the origin of theca cell progenitors and the ovarian lymphatic vasculature (Kinnear et al., 2020), as shown in mice (Liu et al., 2015, Svingen et al., 2012).

During fetal development, the intermediate mesoderm gives rise to both bipotential gonads and adjacent mesonephros structures. The mesonephros initially functions to produce urine, containing glomeruli, mesonephric tubules, the Wolffian duct (or nephric duct or mesonephric duct) and the Müllerian duct (or paramesonephric duct) (Belle et al., 2017, Sainio, 2003, Woolf et al., 2003). In humans, the mesonephri emerge approximately at 3 weeks post fertilization (WPF), prior to sex determination, it produces urine from 6-10WPF and is fully regressed or transformed by 16WPF (Belle et al., 2017, Ludwig and Landmann, 2005, Sainio, 2003, Woolf et al., 2003). In males, the mesonephric tubules transform into the epididymis, the Wolffian duct into the vas deferens and the Müllerian duct regresses. In females, the Müllerian duct develops into the fallopian tube (plus the uterus and part of the vagina), while the Wolffian duct and mesonephric tubules regress (Belle et al., 2017, Sainio, 2003). It remains a matter of debate whether the mesonephros contributes cells to the rete region of the human female and male gonads, however mesonephric cells have been reported to contribute with myoid cells, fibroblasts and endothelial cells to the testis (Sainio, 2003).

Studies in mice have described cells in the rete testis as having both mesonephric and Sertoli-like characteristics (Joseph et al., 2009, Malolina and Kulibin, 2019). It has thus been suggested that the rete network consists of cell types from both origins, making it difficult to identify the molecular markers that define this region. Recently, Kulibin and Malolina investigated rete testis formation in mice between embryonic day (E)11.5 and E16.5 (Kulibin and Malolina, 2020). In mice, rete testis cells express PAX8 (marker of mesonephric tubules), as well as SOX9 and WT1 (Sertoli cell markers). At E13.5, PAX8+/SOX9+/WT1+/AMH-/DMRT1-rete testis cells become canalized and form the rete cords, which connect to PAX8-/SOX9+/WT1+/AMH+/DMRT1+ Sertoli cells in the seminiferous tubes. From E14.5 to E16.5, the AMH+ cells near the rete cords are gradually replaced by PAX8+ rete cells (Kulibin and Malolina, 2020). During this period time, this border region is comprised of a mixed population of PAX8+/DMRT1+ and PAX8+/DMRT1-cells (Kulibin and Malolina, 2020, Major et al., 2021). In mice, single cell transcriptomics has been applied to investigate the developmental stages and cellular trajectories during male gametogenesis (Chen et al., 2018), but the molecular identity of the rete testis was not explored. Similarly, single cell transcriptomics in humans has been used to study testis development and gametogenesis during fetal and adult life (Fan et al., 2021, Guo et al., 2018, Guo et al., 2021, Li et al., 2017, Shami et al., 2020, Sohni et al., 2019, Wang et al., 2018). However, the molecular signatures of the rete testis have not been inferred and differences with the mesonephros remain to be determined.

In this study, we have performed single cell transcriptomics on first trimester (1T) and second trimester (2T) human fetal male and female gonads, focusing on the development of the rete region. In addition, we performed single cell transcriptomics on 1T and 2T male mesonephros/epididymis. We have identified the molecular signature of individual cell types in both fetal gonads and mesonephros/epididymis, with a focus on the characterization of the cells in the rete region.

## RESULTS

### Human fetal rete testis express PAX8 and KRT19 and can contain germ cells

To investigate rete testis development, we have collected human fetal male gonads and mesonephros/epididymis from both 1T and 2T. According to previous studies, PAX8 and KRT19 were specifically expressed in human adult rete testis (Bremmer et al., 2015, Tong et al., 2011), therefore we first confirmed the expression of PAX8 and KRT19 in the human rete testis in the 1T and 2T (Figure 1A, 1B). At 8WPF, the rete testis, expressing PAX8 and KRT19, was not properly organized in tubules, in contrast to the well-organized AMH+ Sertoli-cells in the seminiferous tubes (Figure 1A, 1B). Surprisingly, both DDX4+ germ cells (Figure 1A) and POU5F1+ germ cells (Figure 1B) were observed in the rete testis, suggesting that the rete testis either has a gonadal origin or can attract germ cells. Interestingly, at 8WPF we observed a striking PAX8+KRT19+ tubule with a central lumen inside of the testis in the region where in the future efferent ducts of the epididymis will form (Figure 1A, 1B). Furthermore, at 8WPF we noted a small number of germ cells at the basal part of the tubule (Figure 1B) as well as under the surface epithelial cells of the testis (Figure 1A, 1B). At 17WPF, the PAX8+KRT19+ rete testis formed a well-defined network of tubules that connected with the seminiferous tubes (containing the AMH+ Sertoli cells) and still contained many DDX4+ germ cells (Figure 1A). Regarding the mesonephros, at 8WPF, part of the outer layer of the corpuscle (particularly adjacent to the urinary pole), the mesonephric tubules and the Wolffian duct were PAX8+KRT19+ as well as the efferent tubules and the tubules in the head of the epididymis at 17WPF (Figure 1A, 1B). Finally, the surface epithelial cells at 8WPF and 17WPF expressed low levels of PAX8 and high levels of KRT19 (Figure 1A, 1B).

**Figure 1.**
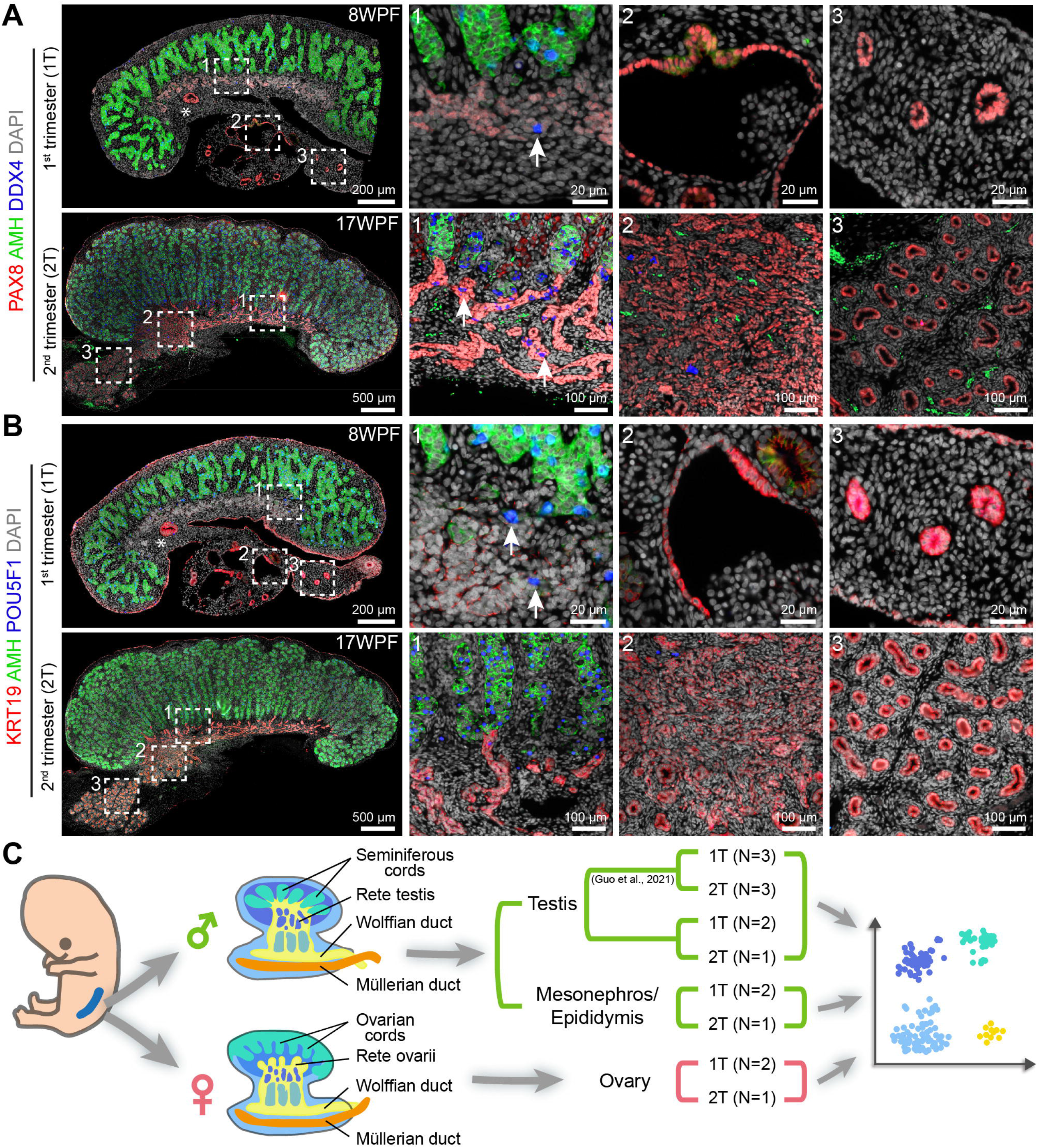
PAX8 and KRT19 expression in the fetal male testes and study set-up. **(A)** Immunofluorescence for PAX8, AMH and DDX4 in first trimester (top) and second trimester testes (bottom). White dashed boxes in the left image indicate selected magnified regions on the right. White asterisk in top left panel indicates PAX8+ tubule in testis. White arrows indicate germ cells in the rete testis. Scale bars are 200 μm in the top left image, 20 μm in the top magnified images, 500 μm in the bottom left image and 100 μm in the bottom magnified images. **(B)** Immunofluorescence for KRT19, AMH and POU5F1 in first trimester (top) and second trimester testes (bottom). White asterisk in top left panel indicates KRT19+ tubule in testis. White dashed boxes in the left image indicate selected magnified regions on the right. White arrows indicate germ cells in the rete testis. Scale bars are 200 μm in the top left image, 20 μm in the top magnified images, 500 μm in the bottom left image and 100 μm in the bottom magnified images. **(C)** Study set-up.

### Identification of rete testis cells in human fetal testes

To define the molecular signature of the cells of the rete testis, we have isolated fetal male gonads and mesonephros/epididymis as well as fetal female gonads and performed single cell transcriptomics (Figure 1C). For a robust identification of the rete testis cells, we have merged our male gonad dataset from 3 donors (9WPF, 9WPF and 18WPF) with a recently published male gonad dataset containing several donors with complementary developmental ages (6, 7, 8, 12, 15, 16WPF) (Guo et al., 2021), as that study did not infer on rete testis identity.

We analyzed a total of 35228 cells (9634 from our dataset and 25594 from Guo and colleagues (Guo et al., 2021)) that passed our quality control. After correcting for batch effects, 19 distinct clusters (mCL0 – mCL18) from 9 donors were identified and all cells were visualized on a two-dimensional plot using uniform manifold approximation and projection (UMAP) algorithm (Figure 2A and S1A). Thereafter, the differentially expressed genes (DEGs) for each cluster were calculated (Table S1). According to cell identity annotations from Guo and colleagues (Guo et al., 2021) and known marker genes, we have confirmed that mCL0 and mCL2 were stromal cells *(NR2F2+TCF21+PDGFRA+),* mCL5 were proliferating stromal cells *(NR2F2+TOP2A+MKI67+)* mCL3 and mCL4 were stromal precursor cells from 6-7WPF *(NR2F2+TCF21+PDGFRA+),* mCL15 were germ cells *(POU5F1+* and/or *DDX4+),* mCL9 were smooth muscle cells *(RGS5+),* mCL6 and mCL13 were endothelial cells *(PECAM1*+), mCL11 were immune cells *(CD53+)* mCL1 and mCL8 were Sertoli cells *(AMH+SOX9+),* mCL10 were Sertoli precursor cells from 6-7WPF *(AMH+SOX9+),* mCL14 were Sertoli interstitial precursor cells *(AMH+SOX9+NR2F2+),* mCL12 were Leydig cells *(CYP17A1+INSL3+),* mCL17 and mCL18 were red blood cells *(HBA1+)* and mCL16 were nephric cells showing expression of podocyte markers *NPHS2, PTPRO* and *CLIC5* (Figure 2A and S1B) (Guo et al., 2021, Hochane et al., 2019). Interestingly, rete testis epithelial cell markers *PAX8* and *KRT19* were mainly expressed in cells from mCL7 (Figure 2B).

**Figure 2.**
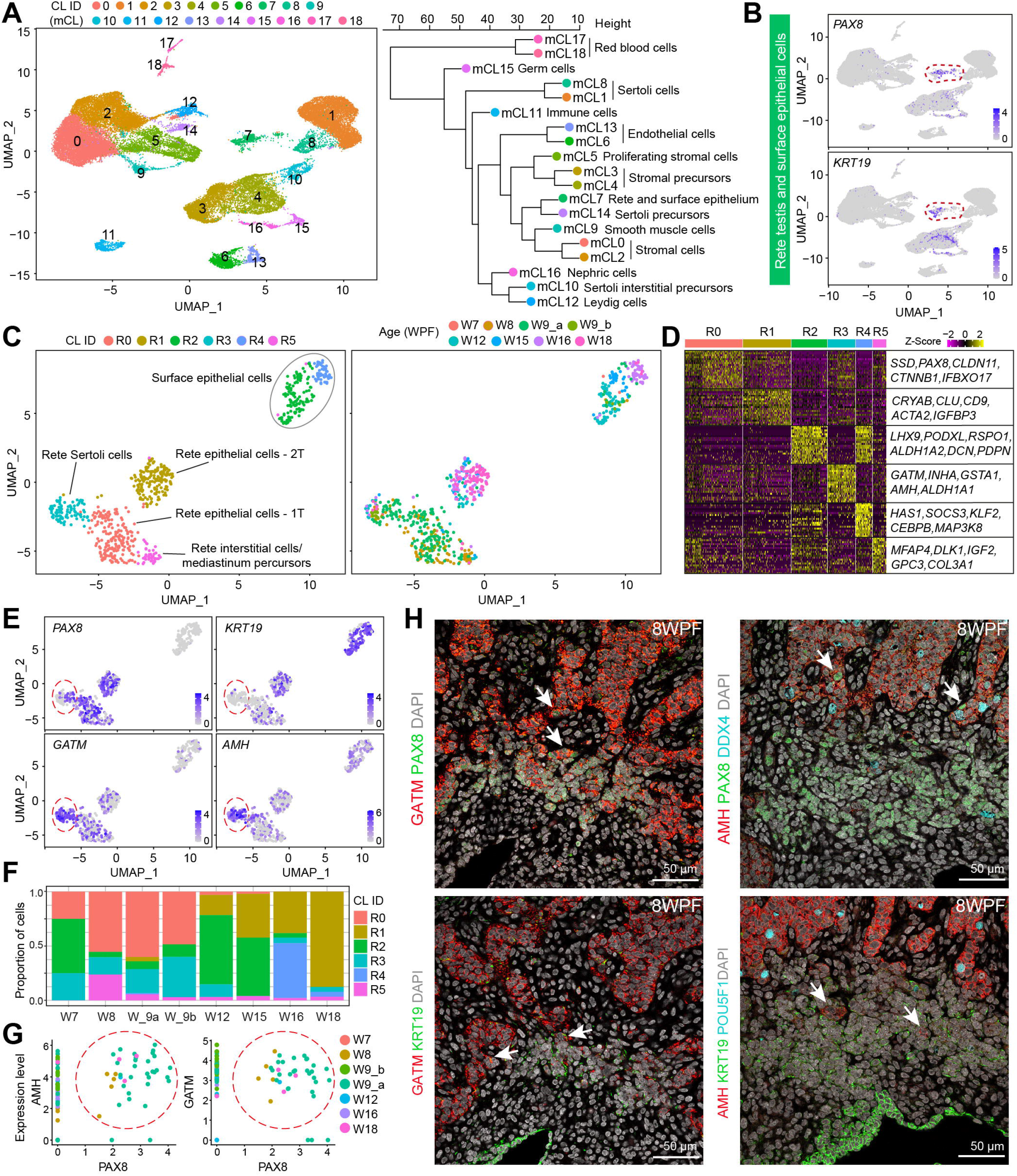
Molecular characterization of cell types in the human fetal rete testis. **(A)** Uniform manifold approximation and projection (UMAP) plot showing cell clusters (mCL) obtained for the human fetal testis (left panel), and hierarchical clustering with the cell cluster identity (right panel). **(B)** UMAP plot showing expression of *PAX8* and *KRT19.* **(C)** UMAP plot showing sub-clustering of mCL7 (left panel) and depicting the developmental age/donor (right panel). **(D)** Heatmap showing the scaled average expression (Z-score) of the top 20 differentially expressed genes (DEGs, ranked by average Log2 fold change) per subcluster. **(E)** UMAP plot of sub-clustering of mCL7 showing expression of *PAX8, KRT19, GATM* and *AMH.* Red dashed circles indicate sub-cluster R3. **(F)** Bar plot showing the proportion of cells from each sub-cluster per donor. **(G)** Scatter plot showing the correlation between expression of *PAX8* and *AMH* (left) and *PAX8* and *GATM* (right) in cells of subcluster R3. Red dashed circles indicate double positive cells. **(H)** Immunofluorescence for GATM and PAX8 (top left), GATM and KRT19 (bottom left), AMH, PAX8 and DDX4 (top right) and AMH, KRT19 and POU5F1 (bottom right) in testis at 8WPF. White arrows indicate cells that are either PAX8+GATM+ or PAX8+AMH+ at the border region between rete testis and seminiferous tubes. Scale bars are 50 μm.

### Transcriptional analysis of fetal male rete epithelial cells

To further explore the cell features of mCL7, we extracted the 679 cells from mCL7 and after sub-clustering we obtained 6 sub-clusters (R0 to R5) (Figure 2C). We next calculated the DEGs for each sub-cluster and plotted a heatmap of top 20 DEGs per subcluster (Figure 2D). We observed that the top 20 DEGs of R2, including *PODXL, ALDH1A2* and *PDPN* were also highly expressed in R4 (Figure 2D and S2A). Validation by immunofluorescence for PODXL, ALDH1A2 and PDPN on the fetal testes showed strong expression on the surface epithelial cells, in addition to high levels of expression of PDPN in germ cells. This strongly suggested that the cells in R2 and R4 corresponded to surface epithelial cells of the testis of 7-15WPF and 16-18WPF, respectively (Figure S2B).

The DEGs of R5, including *DLK1, IGF2, GPC3* and *COL3A1* (Figure 2D), suggested that cells in R5 could correspond to interstitial cells in the rete testis, perhaps the precursors of the mediastinum, as these cells also express low levels of *PAX8* and *KRT19* (Figure 2E).

The high levels of expression of *PAX8* and differential levels of *KRT19* (low in R0 and high in R1) indicated that R0 and R1 represented cells of rete epithelial cells from 1T and 2T, respectively (Figure 2E). Interestingly, the top 20 DEGs of R0 and R1 differed considerably (Figure 2D), suggesting strong developmental dynamics in the rete epithelium between 1T and 2T.

Intriguingly, the top 20 DEGs of R3 included known Sertoli cell markers, such as *AMH, GATM, INHA, GSTA1* and *ALDH1A1* (Figure 2D, 2E and Table S1). Similar to the cells in R0, the cells in R3 were mainly observed in 1T testes (Figure 2F). As we noticed that cells in R3 *(AMH+GATM+)* contained both *PAX8+* and *PAX8-* cells (Figure 2E, 2G), we next investigated the localization of these two populations by immunofluorescence. We observed that AMH+PAX8+ cells or GATM+PAX8+ cells were present at the developing interface between the rete testis and the seminiferous tubes in 8WPF testis (Figure 2H).

Previously, *WT1* and *NR5A1* were detected in Sertoli cells and the rete region (Hanley et al., 1999), whereas *SOX9* was detected only in (seemingly) Sertoli cells during human testis development (Hanley et al., 2000). Moreover, *Dmrt1, Wt1, Sox9* and *Nr5a1* have been described in both the rete testis and Sertoli cells in mice, whereas rete cells could be distinguished from Sertoli cells by the combined expression of *Cdh1* and *Krt8* (Major et al., 2021). Cells in R3 showed specific enrichment for *DMRT1, NR5A1* and *SOX9* as well as high levels of *KRT8* and *WT1*, while *CDH1* was not detected (Figure S2C). It remains to be investigated whether Sertoli cells in the rete testis differentiate from Sertoli cells in the seminiferous tubes or from the rete testis epithelial cells.

### Single-cell transcriptomics of human fetal male mesonephros and epididymis

Next, we investigated the sex-specific transition from male mesonephros into epididymis by performing single cell transcriptomics and analyzing the obtained datasets from human fetal mesonephros/epididymis of 3 different donors (7WPF, 12WPF, 17WPF) (Figure 1C, 3A). Using the same thresholds for quality control as for the testis, we retained 8707 cells and obtained 16 different clusters (meCL0 – meCL15) by UMAP analysis (Figure 3A). The analysis of the DEGs calculated for each meCL (Table S2) as well as expression of known marker genes, suggested that clusters meCL6, meCL7 and meCL11 showed similarities with nephric epithelial cells *(PAX2+LHX1+)* (Boualia et al., 2013, Hochane et al., 2019, Takasato et al., 2014), meCL15 were endothelial cells *(PECAM1+)* meCL4 were Sertoli cells *(AMH+)* from 7WPF, meCL9 were Sertoli cells *(AMH+)* probably from contamination with 17WPF testis while isolating the epididymis, and meCL14 showed cells with similarities with nephric podocytes *(PODXL+CLIC5+)* (Hochane et al., 2019) (Figure 3B). Using immunofluorescence, we confirmed the presence of PODXL+ podocytes as well as PODXL+ cells in the outer layer of the corpuscle, the vascular endothelial cells, the surface epithelium of the mesonephros and the Mullerian duct, but not in the mesonephric tubules at 8WPF (Figure 3C).

**Figure 3.**
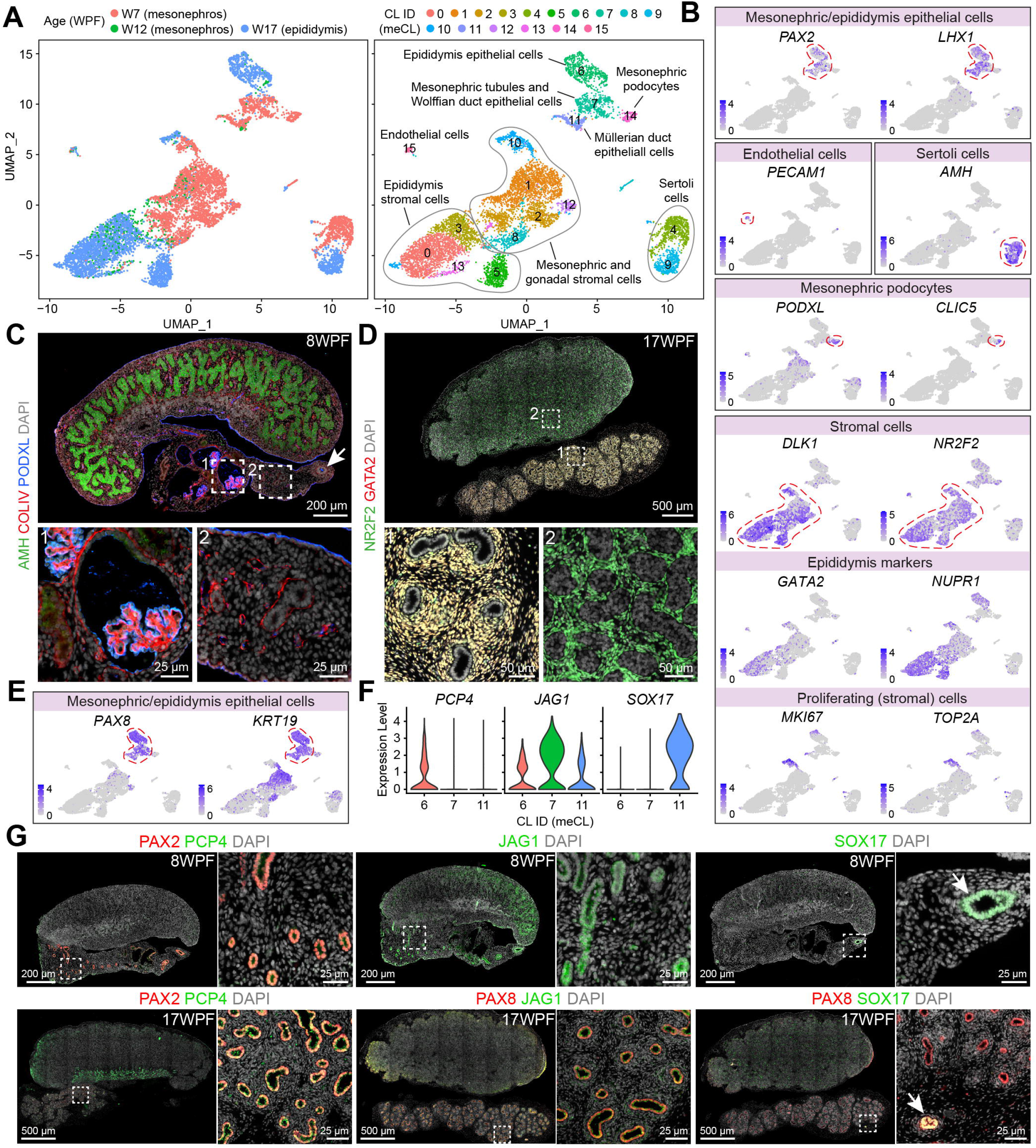
Molecular characterization of cell types in human fetal male mesonephros and epididymis. **(A)** Uniform manifold approximation and projection (UMAP) plot showing the different donors (left panel) and the cell cluster identity (meCL) (right panel) obtained for the human fetal mesonephros and epididymis (left panel). **(B)** UMAP plot showing expression of selected markers. Red dashed circles indicate cell populations with specific marker expression. **(C)** Immunofluorescence for AMH, COLIV and PODXL in testis and mesonephros at 8WPF. White dashed boxes in the top image indicate selected magnified regions on the bottom. White arrow indicates the Müllerian duct. Scale bars are 200 μm in the top and 25 μm in bottom panels. **(D)** Immunofluorescence for NR2F2 and GATA2 in testis and epididymis at 17WPF. White dashed boxes in the top image indicate selected magnified regions on the bottom. Scale bars are 500 μm in the top and 50 μm in bottom panels. **(E)** UMAP plot showing expression of *PAX8* and *KRT19.* Red dashed lines indicate epithelial cell clusters. **(F)** Violin plots showing expressions of *PCP4, JAG1* and *SOX17* in epithelial cell clusters (meCL6, meCL7 and meCL11). **(G)** Immunofluorescence for PAX2 and PCP4 (right panels), JAG1 (middle panels) and SOX17 (right panels) in testis and mesonephos at 8WPF (top row); and for PAX2 and PCP4 (right panels), PAX8 and JAG1 (middle panels) and PAX8 and SOX17 (right panels) in testis and epididymis at 17WPF (bottom row). White dashed boxes indicate the selected magnified region for each staining. White arrows indicate the (regressing) Müllerian duct. Scale bars are 500 μm in the overview and 25 μm in the magnification.

The stromal compartment was represented by 9 clusters (meCL0, meCL1, meCL2, meCL3, meCL5, meCL8, meCL10, meCL12 and meCL13), expressing high levels of *DLK1* and *NR2F2* (Figure 3B). Differentiating the mesenchyme from the male mesonephros and the epididymis at 17WPF, we noticed high levels of *GATA2* and *NUPR1* (Figure 3B). We were able to validate the co-expression of GATA2 and NR2F2 in the mesenchyme of the epididymis at 17WPF by immunofluorescence (Figure 3D).

Next, we focused on the epithelial cells in meCL6, meCL7 and meCL11 and, as expected from immunofluorescence results (Figure 1A, 1B), they all showed high levels of expression of *PAX8* and *KRT19* (Figure 3E). In addition, these cells also showed high expression of *KRT8* and moderate expression of *SOX9* (Figure S2D). However, *CDH1* and *WT1* were only faintly detected in epithelial cells in 1T (Figure S2D). To discriminate between the 3 epithelial clusters, we selected *PCP4, JAG1* and *SOX17* (Figure 3F and S2E) from the obtained DEGs (Table S2) for validation by immunofluorescence. Interestingly, PCP4 not only colocalized with PAX2 in the tubules of the epididymis at 17WPF, but it proved to be a specific marker of the tubules in rete testis (Figure 3G). The expression of PCP4 in the rete testis is in agreement with *PCP4* being the top DEG of cluster mCL7, identified as rete testis and testis surface epithelium (Table S1). JAG1 was highly expressed in the mesonephric tubules at 8WPF and moderately expressed in the tubules of the epididymis at 17WPF (Figure 3G).

Recently, SOX17 has been described as a marker of epithelial cells in the human adult fallopian tube, that derive from the Müllerian duct, and it is suggested that SOX17 and PAX8 could even drive differentiation of the fallopian tube epithelium (Dinh et al., 2021). In our dataset from mesonephros/epididymis, *SOX17* was specifically expressed in meCL11 (Figure 3F) and immunofluorescence SOX17 revealed the Müllerian duct in the male mesonephros at 8WPF and the regressing Müllerian duct in the epididymis of 17WPF testis (Figure 3G). In addition, we also confirmed expression of SOX17 in the Müllerian duct in the female mesonephros and female germ cells at 8WPF (Figure S2F).

### Comparison between fetal male rete and mesonephric/epididymis epithelial cells

In our analysis, we have identified rete epithelial cells in the testis (R0, R1 and R3) and male mesonephric/epididymis epithelial cells (meCL6, meCL7, meCL11), all expressing *PAX8* and *KRT19.* To directly compare the molecular signature of these two sets of epithelial cells, we merged the cells of R0, R1 and R3 and compared the expression profile of that group (R0+R1+R3) to a group formed by an equal number of cells randomly selected from the rest of the testis dataset (Figure S2G). We performed this cell reduction by random cell selection. We visualized the gene expression differences in a Volcano plot (adjusted p-value <0.05 and average log2 transformed fold change >0.5) (Figure 4A) and identified 453 DEGs upregulated in the group of rete epithelial cells (Table S3).

**Figure 4.**
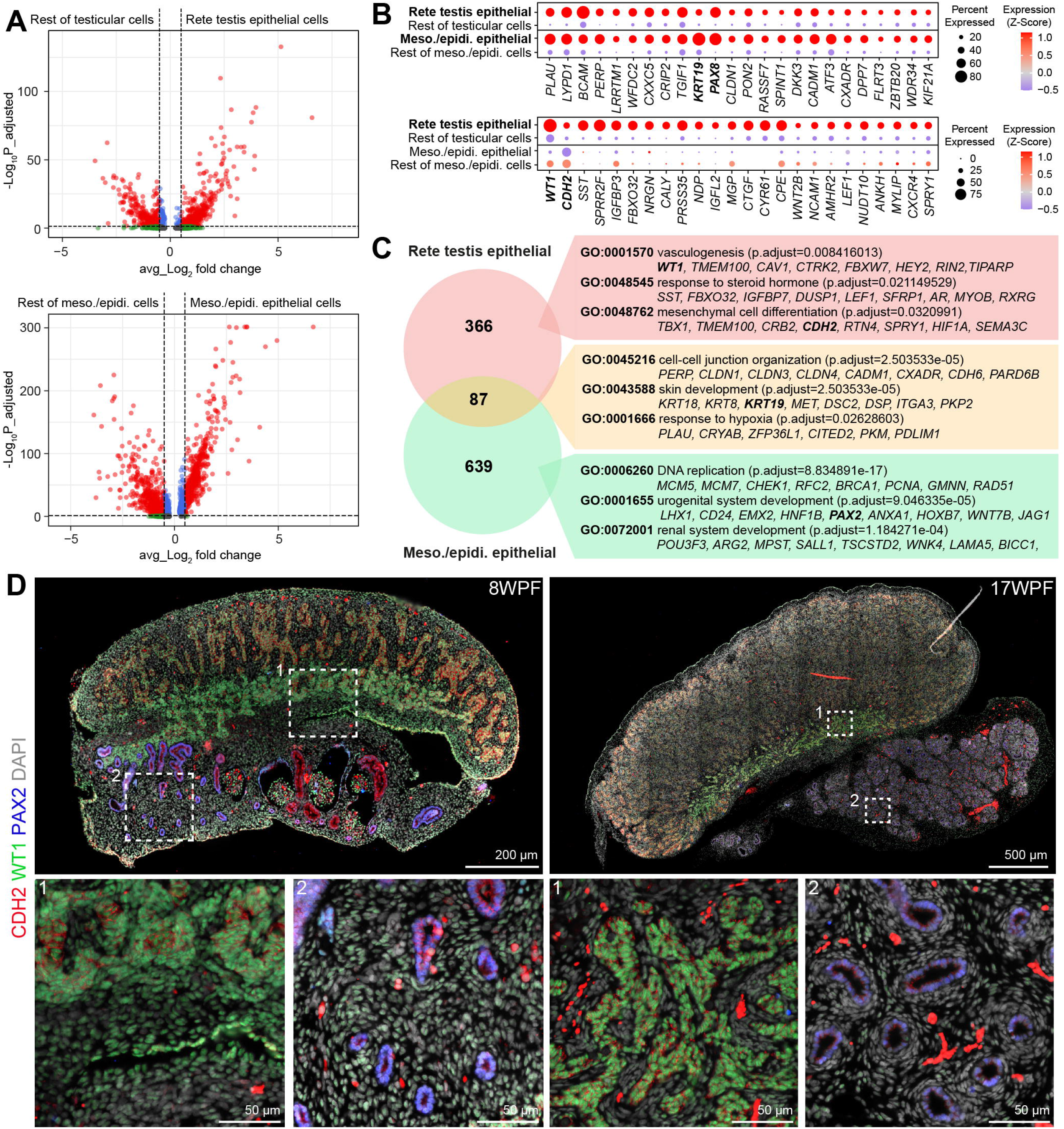
Common DEGs between male rete and mesonephric/epididymis epithelial cells. **(A)** Volcano plots showing differential expression between male rete epithelial cells and the rest of the testicular cells (top panel) and differential expression between male mesonephros and epididymis epithelial cells and the rest of cells in the mesonephros and epididymis (bottom panel). **(B)** Dot plots showing the scaled expression (Z-score) of selected genes differentially upregulated in both male rete and mesonephric/epididymis epithelial cells in the populations of interest (top) and showing the scaled expression (Z-score) of selected genes differentially upregulated only in male rete epithelial cells in the populations of interest (bottom). **(C)** Venn diagram showing the intersection between the genes significantly upregulated in the male rete epithelial cells and the genes significantly upregulated in the mesonephros and epididymis epithelial cells. On the right part are representative GO terms (biological processes) and representative genes. **(D)** Immunofluorescence for CDH2, WT1 and PAX2 in testis and mesonephros at 8WPF (left) and testis and epididymis at 17WPF (right). White dashed boxes in top image indicate the selected magnified images in the bottom. Scale bars are 200 μm in 8WPF and 500 μm in 17WPF overview images and 50 μm in the magnified images.

We then merged the cells in meCL6, meCL7, meCL11 and compared the expression profile of that group (meCL6+meCL7+meCL11) to a group formed by an equal number of cells randomly selected from the rest of the mesonephros/epididymis dataset (Figure S2H). We also visualized the gene expression differences in a Volcano plot (adjusted p-value <0.05 and average log2 transformed fold change >0.5) (Figure 4A) and identified 726 DEGs upregulated in the group of mesonephros/epididymis epithelial cells (Table S3).

The intersection between the DEGs upregulated in the rete testis epithelial cells (453 DEGs) and the DEGs upregulated in mesonephros/epididymis epithelial cells (726 DEGs) revealed 87 common genes (Table S3), including *KRT19* and *PAX8* (Figure 4B). The Gene Ontology (GO) terms for biological processes that were enriched using the 87 common genes mainly associated with epithelial cell features, such as “cell-cell junction organization” and “skin development” (Figure 4C and Table S4 for complete list). Moreover, we observed genes, such as *PLAU, CRYAB, ZFP36L1,* associated with GO term “response to hypoxia” (Figure 4C), in agreement with several studies that have described that hypoxic conditions are important for mesonephros and male gonadal development in quail and mouse, respectively (Kirschner et al., 2017, Nanka et al., 2006).

From the unique upregulated DEGs in the rete testis epithelial cells (366 DEGs), including *WT1* and *CDH2* (Figure 4B), we observed GO terms for “vasculogenesis”, “response to steroid hormone” and “mesenchymal cell differentiation” (Figure 4C and Table S4 for complete list); whereas from the unique upregulated DEGs in the mesonephros/epididymis epithelial cells (639 DEGs) we observed GO terms linked with “DNA replication”, “urogenital system development” and “renal system development” (Figure 4B and Table S4 for complete list). Using immunofluorescence, we were able to validate the expression of WT1 and CDH2 in the epithelial cells of the rete testis, but not in the epithelial cells of the PAX2+ mesonephros/epididymis epithelial cells (Figure 4D).

### Identification of rete ovarii cells in human fetal ovary

Next, we investigated the molecular signature of the cells in the female gonad (Figure 1C, 5A) to determine differences and similarities of rete cells between the sexes. For this, we collected fetal human female gonads from 1T (2 donors from 9 WPF) and 2T (one donor from 16 WPF) and performed single cell transcriptomics.

**Figure 5.**
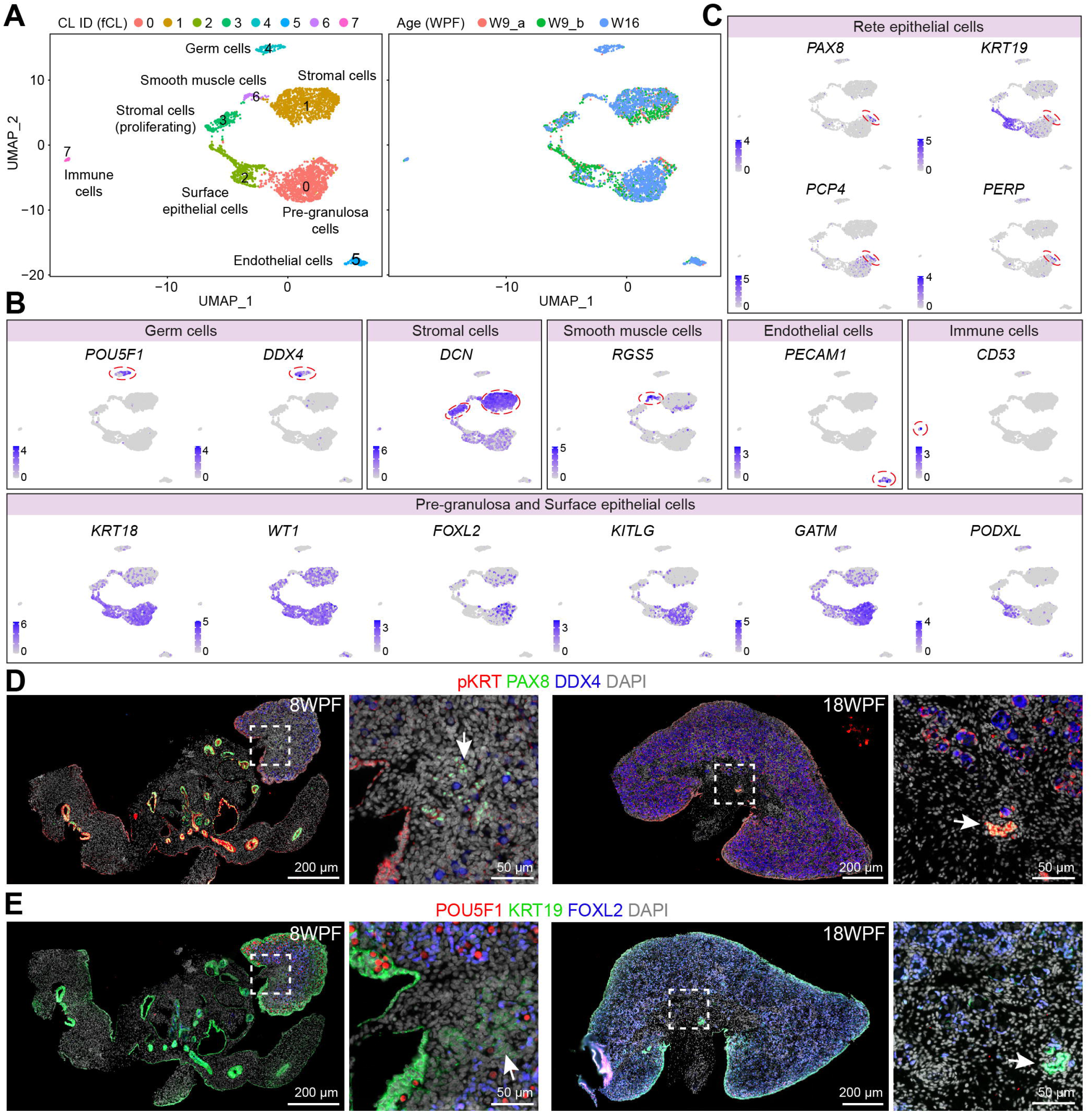
Molecular characterization of cell types in human fetal ovary. **(A)** Uniform manifold approximation and projection (UMAP) plot showing the cell cluster identity (fCL) (right panel) and the developmental age/different donors (left panel) obtained for the human fetal ovary. **(B)** UMAP plot showing expression of selected markers for major cell types in fetal ovaries. Red dashed circles indicate cell populations with specific marker expression. **(C)** UMAP plot showing expression of selected markers of the rete testis. Red dashed circles indicate cell populations with specific marker expression. **(D)** Immunofluorescence for pan (p)KRT, PAX8 and DDX4 in ovaries and mesonephros at 8WPF and ovary at18WPF. White dashed boxes in the overview images indicate the magnified region to the right. White arrows indicate cells equivalent to the rete testis in the ovary, the rete ovarii. Scale bars are 200 μm in the overview and 50 μm in the magnification. **(E)** Immunofluorescence for POU5F1, KRT19 and FOXL2 in ovaries and mesonephros at 8WPF and ovary at18WPF. White dashed box in the overview image indicates the magnified region to the right. White arrow indicates cells equivalent to the rete testis in the ovary, the rete ovarii. Scale bars are 200 μm in the overview and 50 μm in the magnification.

In total, 3509 female gonadal cells passed our quality control thresholds. Using a Seurat-based anlysis, we identified 8 distinct clusters (fCL0 - fCL7) (Figure 5A) and the DEGs for each cluster were calculated (Table S5). Based on the DEGs and several known marker genes (Li et al., 2017), we were able to identify the cells in fCL4 as fetal germ cells *(POU5F1+* or *DDX4*+), cells in fCL1 and fCL3 were stromal cells (*DCN+*) and fCL3 contained proliferating stromal cells *(DCN+MKI67+TOP2A+),* cells in fCL6 were smooth muscle cells (*RGS5*+), cells in fCL5 were endothelial cells (*PECAM1*+), cells in fCL7 were immune cells (*CD53*+), cells in fCL2 were surface epithelial cells *(PODXL+KRT18+)* and cells in fCL0 were pre-granulosa cells *(FOXL2+KITLG+KRT18+)* (Figure 5B and S3A). We validated the expression of PODXL in the surface epithelium cells in 1T and 2T gonads (Figure S3B) as in males (Figure S2B) and show that WT1 is expressed in the somatic compartment of the ovary, but not in the fetal germ cells (Figure S3C).

When investigating the expression of rete testis markers in the fetal ovary dataset, we noticed high levels of expression of *PAX8, PCP4* and *KRT19* in a group of cells from fCL0 (pre-granulosa cells), that could correspond to rete ovarii cells (Figure 5C). Interestingly, as in males, the female surface epithelium cells (fCL2) were *KRT19+PAX8-* (Figure 2E, 5C). Surprisingly, immunofluorescence for PAX8 and KRT19 revealed a core of rete ovarii cells at 8WPF, that in contrast to males in 1T did not organize in tubules (Figure 5D, 5E). Moreover, as in 1T males, PAX8 and KRT19 were also strongly expressed in the mesonephros tubules, Wolffian duct and Müllerian duct in females at 8WPF (Figure 5D, 5E). Interestingly, we were able to identify the remnants of a narrow epithelial structure in the rete ovarii, that was both PAX8+ and KRT19+ in the ovary at 18WPF (Figure 5D, 5E).

### Comparison of fetal rete epithelial cells between the sexes

To further investigate the identity the *KRT19+PAX8+PCP4+* cells identified in the dataset of the female gonads, we have sub-clustered fCL0 and obtained 5 sub-clusters (G0 to G4) (Figure S3D). We next analyzed the distribution of the top 50 most variably expressed genes in the 5 sub-clusters (Figure S3E). From this, we identified sub-cluster G4, with expression of several genes, such as *LYPD1, IGFBP3, PCP4, PLAU, PAX8,* that were previously identified as DEGs in the rete testis epithelial cells (Figure 4B). Therefore, we concluded that the cells from G4 are likely rete ovarii epithelial cells, the female equivalent to the male rete testis epithelial cells, present as a small percentage of cells in the female gonads of 1T (Figure 6A, 6B).

**Figure 6.**
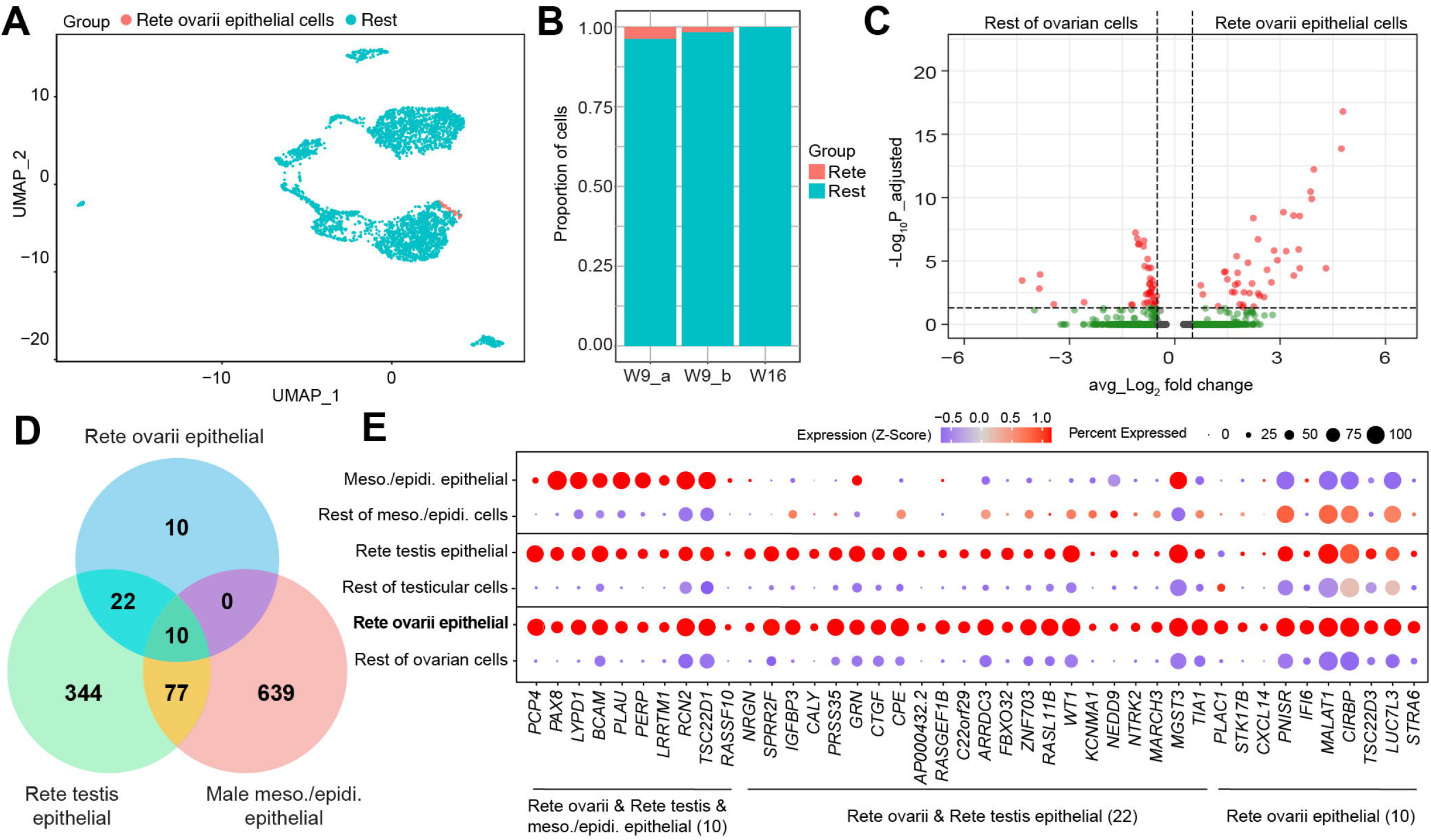
Analysis of the rete ovarii epithelial cells and comparison between sexes. **(A)** Uniform manifold approximation and projection (UMAP) plot showing the identified rete ovarii epithelial cells in the total dataset of fetal ovarian cells. (**B**) Bar plot showing the proportion of rete ovarii epithelial cells per donor. **(C)** Volcano plot showing differential expression between rete ovarii epithelial cells and the rest of the fetal ovarian cells. **(D)** Venn diagram showing the intersection between the genes significantly upregulated in the rete ovarii epithelial cells, the genes significantly upregulated in the male rete epithelial cells and the genes significantly upregulated in the mesonephros and epididymis epithelial cells. **(E)** Dot plots showing the scaled expression (Z-score) of the genes significantly upregulated in the rete ovarii epithelial cells in the other populations of interest.

To compare the molecular signature of the fetal rete epithelial cells between the sexes more thoroughly, we determined the gene expression differences between cells in G4 (39 cells) and a group of 100 cells randomly selected from the rest of the female gonad dataset (Figure S3F). We visualized the gene expression differences in a Volcano plot (adjusted p-value <0.05 and average log2 transformed fold change >0.5) (Figure 6C) and identified 42 DEGs upregulated in the group of the rete ovarii epithelial cells. We then intersected the 42 DEGs with the previously obtained DEGs upregulated in the rete testis epithelial cells (453 DEGs) and the DEGs upregulated in mesonephros/epididymis epithelial cells (726 DEGs) (Figure 6D). We have obtained a set of 10 commonly upregulated genes between all three datasets and 22 commonly upregulated genes between female and male rete epithelial cells, including *SPRR2F, CTGF* and *IGFBP3.* Finally, only 10 genes were specifically upregulated in the rete ovarii epithelial cells (Figure 6D, 6E and Figure S3G), suggesting that the rete ovarii epithelial cells still shares many similarities with the rete testis epithelial cells.

## DISCUSSION

The morphogenesis of the rete testis and the efferent ducts have been often neglected in studies of human gonadal development. However, the rete testis, efferent ducts and Wolffian duct together form a continuous tubular system that plays an essential role in the transport of the sperm cells (de Mello Santos and Hinton, 2019) and hence male fertility. Studies using knock-out mouse models have revealed several genes, such as *Pax2* and *Gata3,* that are associated with defects in mesonephros development; or *Ar* and *Inhba* that are associated with the maintenance of the Wolffian duct (Murashima et al., 2015), but genes affecting the development of the male reproductive tract in humans are not well investigated. By analyzing the molecular make-up of the cells that constitute the human fetal testis, the male mesonephros and the epididymis, we provide an overview of the distinguishing features of the cell types present in these organs at the molecular level.

We have investigated the transcriptomic landscape of human male fetal mesonephros and epididymis and revealed markers for specific structures and cell types. For example, we have shown that GATA2 was expressed in the mesenchyme of the epididymis, SOX17 marked the Müllerian duct epithelial cells and PAX2 marked all epithelial cells of the mesonephros and epididymis. Moreover, it is not surprising that mesonephric and epididymal epithelial cells express known nephric lineage markers, such as *PAX2, PAX8* and *LHX1* (Bouchard et al., 2002, Khoshdel Rad et al., 2020), as the human kidney (or metanephros) develops from an epithelial structure, the ureteric bud, that sprouts from the caudal portion of the Wolffian duct and penetrates a specific region in the intermediate mesoderm, the metanephric mesoderm (Rumballe et al., 2010, Woolf et al., 2003).

Regarding the human rete testis, our analysis has revealed striking similarities between human rete testis epithelial cells and mesonephric/epididymis epithelial cells, including high expression of *PAX8, KRT19, PCP4, PERP, LYPD1* and *BCAM,* perhaps suggesting a common origin. On the other hand, we have identified genes, such as *CDH2* and *WT1* that are common between Sertoli cells and rete testis epithelial cells, but absent from mesonephric/epididymis epithelial cells, suggesting that both the seminiferous tubes and rete testis are undergoing cell sorting and tubule formation, rather than sharing a common origin. As previously reported in mice (Kulibin and Malolina, 2020, Malolina and Kulibin, 2019), we also identified a small population of human cells at the interface between the seminiferous tubes and the rete testis expressing *PAX8,* as well as Sertoli cell markers such as *AMH, SOX9* and *NR5A1.* This suggests that these cells harbored Sertoli-like features, but their origin remains to be investigated. In addition, we have identified common upregulated genes in both male and female rete cells compared to all other cells. These include epithelial specific reactive oxygen damage protector gene *SPRR2F* (Huynh et al., 2020, Yoo et al., 2018), testicular apoptosis regulator gene *IGFBP3* (Lue et al., 2010) and Hippo/YAP1-signalling target gene *CTGF* (Shome et al., 2020).

Surprisingly, we observed the presence of (POU5F1+ and or DDX4+) germ cells in the rete tubules. We suggest that these could have been engulfed during the formation of the rete tubules and it remains unclear during further development whether they enter apoptosis or migrate towards the seminiferous tubes. Although the human and mouse fetal rete testis shared some similarities, such as expression of *PAX8* and *WT1*, one striking difference was the presence of *CDH1* in mouse rete testis from E14.5 onwards (Kulibin and Malolina, 2020), while *CDH1* was absent from human rete testis. Instead, we have detected high levels of CDH2 in both Sertoli cells and rete testis epithelial cells in human, suggesting species-specific differences.

We have identified a population of (PODXL+) surface epithelial cells in both male and female gonads. In human female gonads, both pre-granulosa cells, surface epithelial cells and rete ovarii cells expressed KRT19, but rete ovarii cells could be distinguished by the coexpression of PAX8. It has been recently showed in mice that surface epithelial cells will give rise to pre-granulosa cells that form quiescent follicles, but the pre-granulosa cells that support fertility would have a medullary origin (Niu and Spradling, 2020). Moreover, a recent study on the ontogeny of gonadal supporting cells in mice and non-human primates identified that progenitors of pre-granulosa cells express epithelial genes and cytokeratins (Sasaki et al., 2021). In humans, it remains to be investigated whether surface epithelial cells contribute to the formation of pre-granulosa cells with the capacity to contribute to the pool of functional follicles.

In conclusion, we have analyzed the transcriptome of human fetal gonads from both sexes and male mesonephros/epididymis from 1T and 2T and provided a comprehensive overview of the molecular profiles of rete and mesonephros/epididymis epithelial cells. Notably, we have also identified a small population of rete ovarii cells in 1T showing unexpected similarities to male rete epithelial cells, regarding their gene expression profile. The provided datasets may serve as a valuable resource not only to shed a light on the process of sexual differentiation in humans, but also to provide insights on the genetic foundation underlying the development of the male reproductive ductal system.

## ACKNOWLEDGMENTS

We are very grateful to the staff of the Vrelinghuis, Utrecht and Gynaikon, Rotterdam for their efforts in obtaining the human fetal material as well as the donors that have consented for the use of the material. In addition, we would like to thank D. Cats for helping with data analysis, S.M. Kiełbasa for assistance with the demultiplexing of the samples, M. Bialecka for assistance in the collection of the gonadal samples for sequencing and M. Ferreira for discussions about the human rete testis.

## AUTHOR CONTRIBUTIONS

JT, SMC, XF and SMCDSL conceived and designed the study. SMC, SMCDSL, SH and TVDH collected samples. JT, SMC, SH, TVDH, YWC and XF carried out experiments. JT, XF, HM and IM performed bioinformatic analyses. JT, SMC, XF, IM, YWC, SH, TVDH, HM and SMCDSL analyzed and interpreted data. JT, SMC, XF and SMCDSL wrote the manuscript. All authors wrote, provided critical feedback and approved the final version of the manuscript.

## STAR METHODS

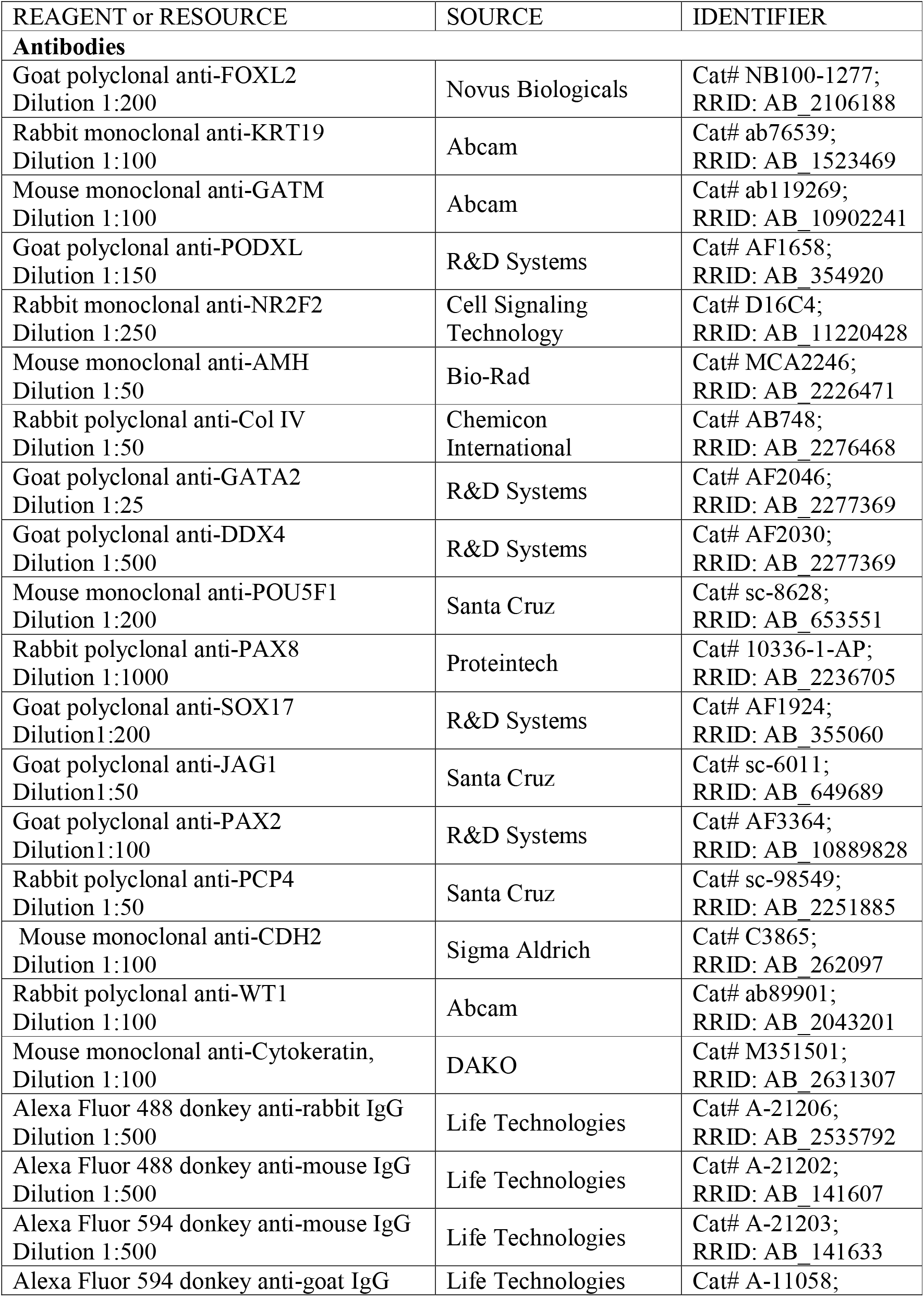
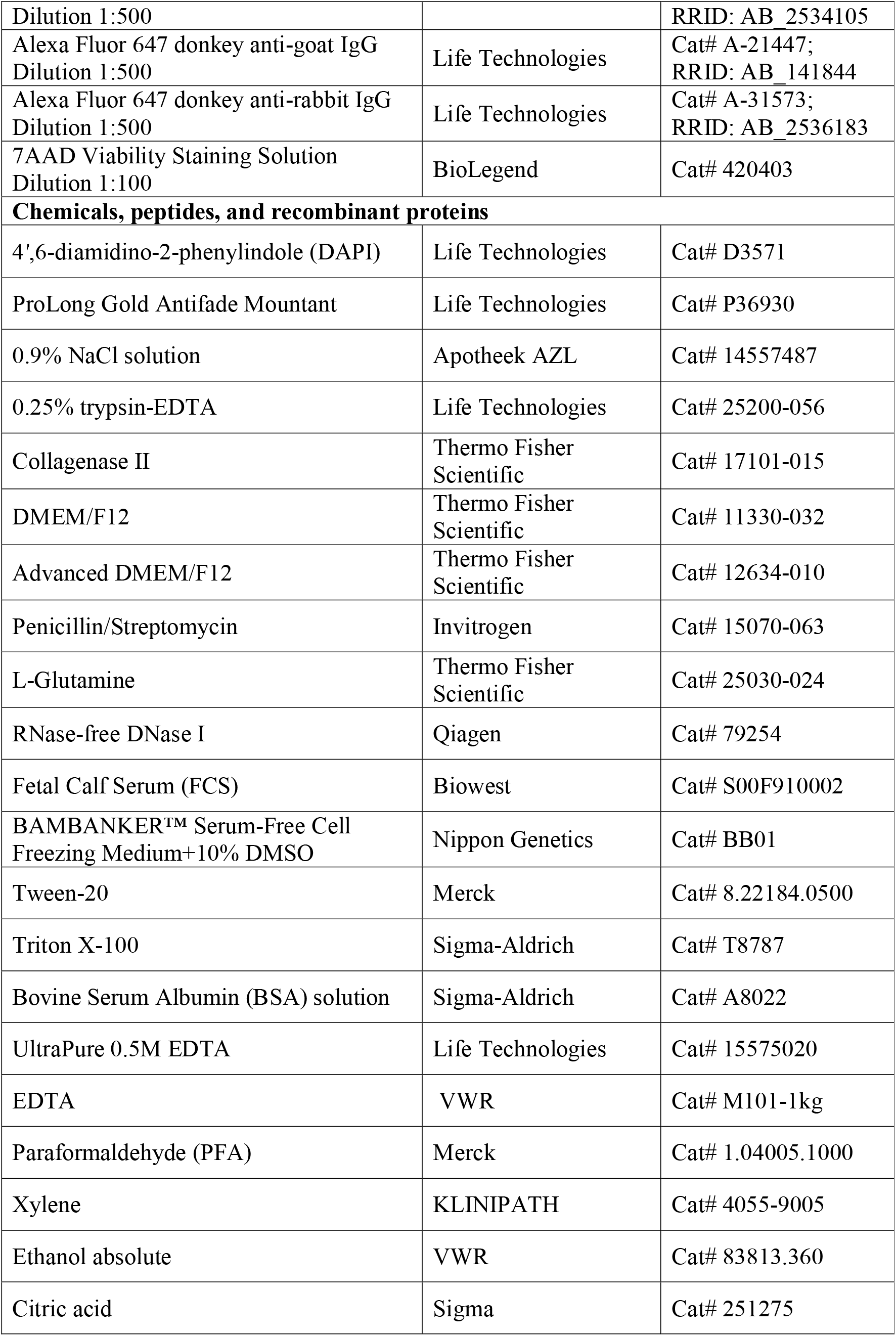
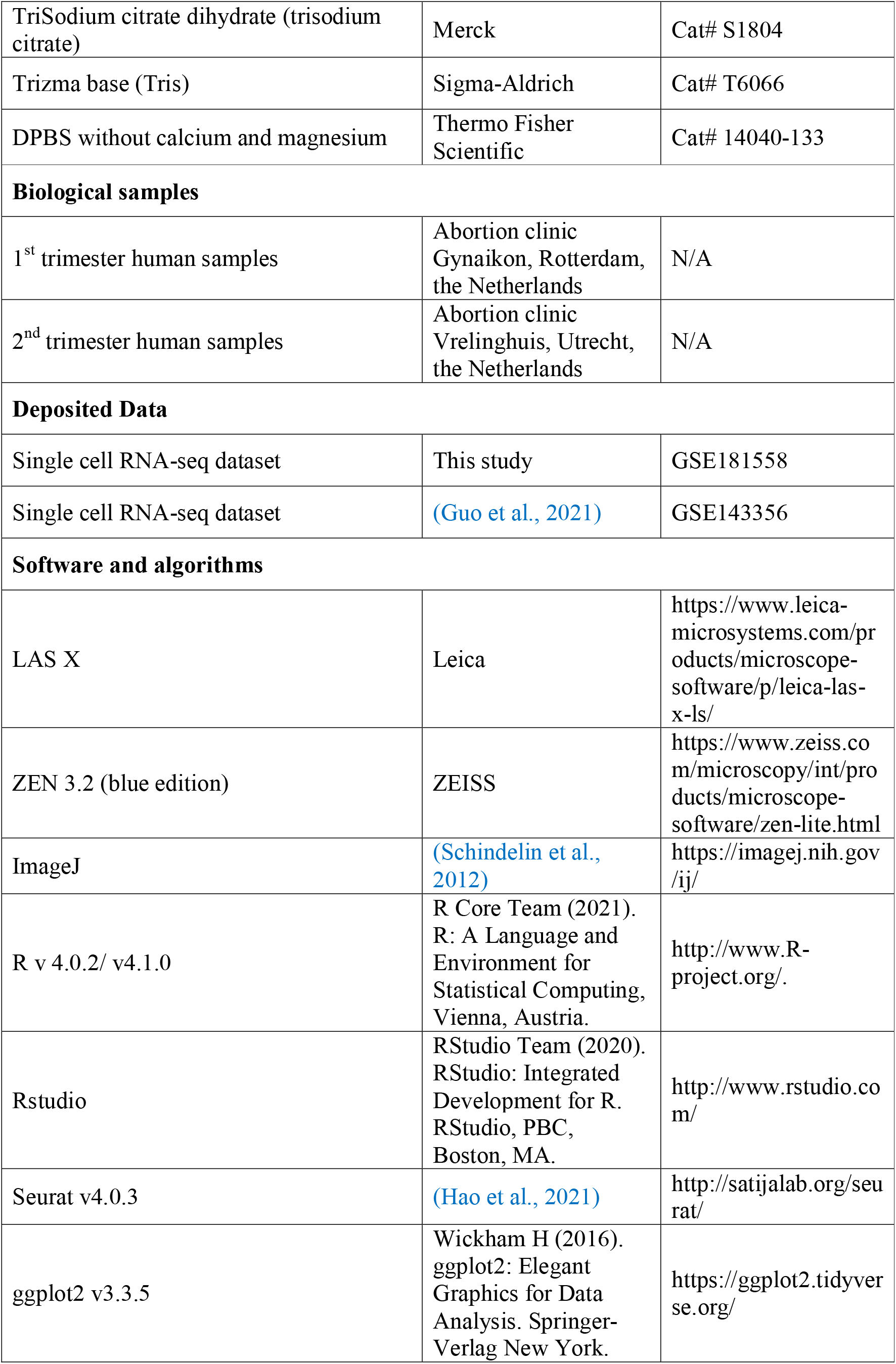
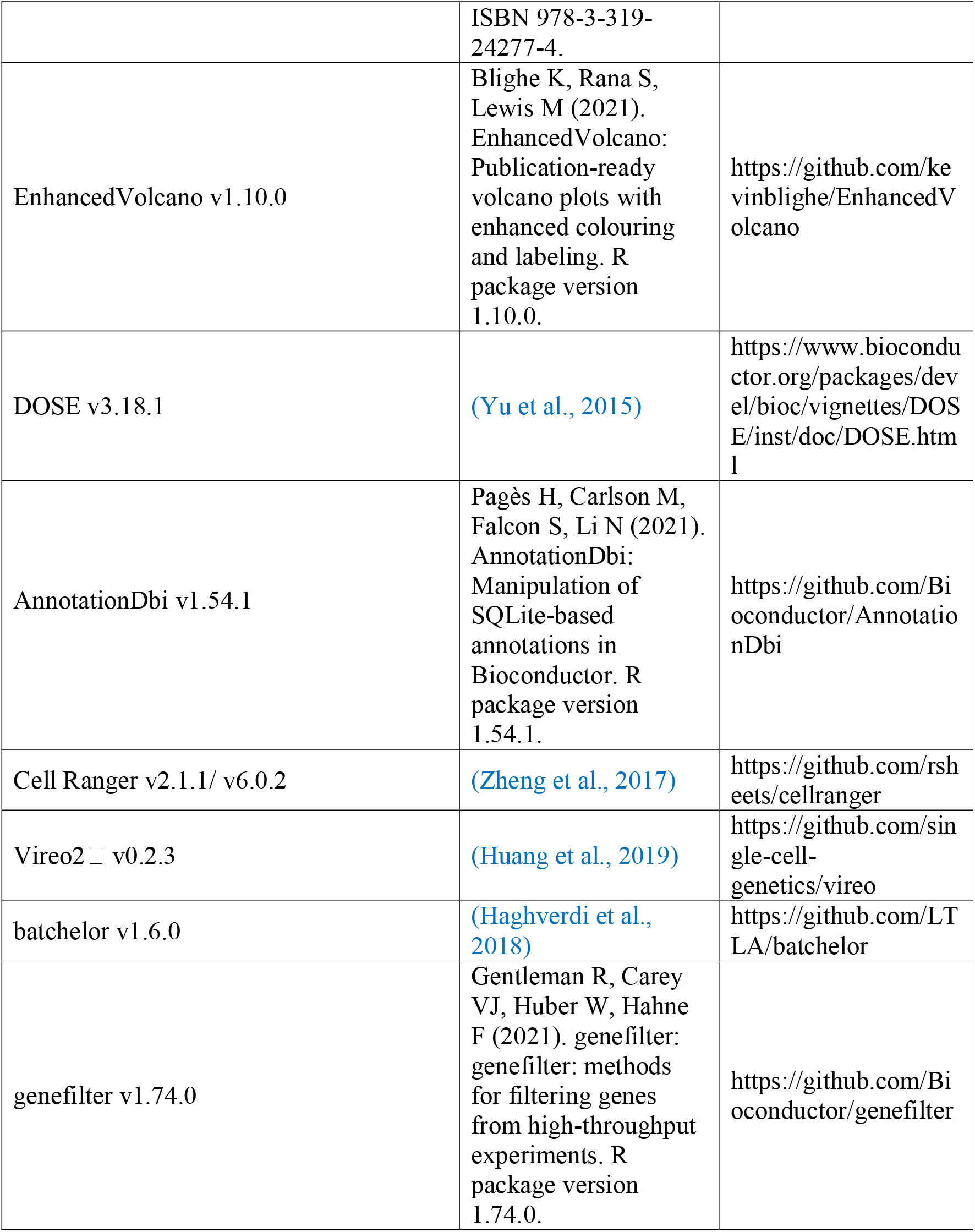
Key Resource Table.

## RESOURCE AVAILABILITY

### Lead contact

Further information and requests for resources and reagents should be directed to and will be fulfilled by the lead contact: Prof. Dr. S. M. Chuva de Sousa Lopes.

### Materials Availability

This study did not generate new unique reagents.

### Data and code availability

The accession number for the RNA sequencing and processed data reported in this paper is Gene expression omnibus (GEO): GSE181558. The code used in the manuscript will be made available for the review process and for publication on github.

## METHOD DETAILS

### Ethical statement and sample collection

All experiments performed in this study were conformed to the Declaration of Helsinki for Medical Research involving Human Subjects and a letter of no objection was obtained from by the Medical Ethical Committee of the Leiden University Medical Center (P08.087). All human fetal material used in this study was obtained with informed consent from the donors, after elective abortions without medical indication.

The developmental age of the fetal samples was estimated in weeks post fertilization (WPF) by ultrasonography. The human fetal gonads, mesonephros and epididymis were dissected in 0.9% NaCl solution for further use.

### Preparation and sequencing of single-cell RNA-Seq libraries

Human fetal gonads, mesonephros and epididymis were prepared for single cell RNA sequencing as previously described (Hochane et al., 2019). Briefly, human fetal gonads, mesonephros and epididymis were incubated overnight at 4°C in an enzyme mix solution (0.1% collagenase II in 0.25% trypsin-EDTA solution). The following day, the samples were centrifuged at 160x*g* for 3 minutes (min) and incubated in Advanced DMEM/F12 containing 1x Penicillin/Streptomycin, 1x L-Glutamine and 27.3IU/ml RNase-free DNase I, at 37°C for 30min to 1 hour. Next, the cell suspension was pipetted up and down for 1min and the enzyme activity was neutralized by adding 10% FCS. The samples were filtered through a 100μm cell strainer (Corning) and centrifuged at 160x*g* for 3min. The single cell suspension was cryopreserved in Bambanker and stored in liquid nitrogen until use.

The single cell suspensions were thawed and resuspended in FACS buffer (1% BSA and 2□mM UltraPure EDTA in DPBS without calcium and magnesium). The cell suspensions were filtered through the FACS tube strainer cap and treated with 7AAD on ice for 3 min. Live cells were sorted on a BD FACS Aria I (BD Biosciences) system, equipped with a 50μm nozzle, 405nm laser, a 695/40A long pass filter and BD FACSDiva 8.0.1 software. Cells were collected in 1% BSA in DMEM/F12 with 1x Penicillin/Streptomycin. The collected live cells were sent to the Leiden Genome Technology Center (LGTC) for library preparation using the 10x Genomics Chromium single cell 3’ Reagent v2 kit (gonads) or 10x Genomics Chromium single cell 3’ Reagent v3 kit (mesonephros/epididymis) according to the manufacturers’ instructions (Zheng et al., 2017) and sequenced.

### Processing and (statistical) analysis of single cell RNA-Seq data

Raw RNA sequencing data of human gonads was initially processed using the Cell Ranger pipeline (v2.1.1) and raw data of human mesonephros/epididymis was processed with Cell Ranger software (v6.0.2). Reads were aligned to the human reference genome (GRCh38). The R package Vireo2 (v0.2.3)□ was applied to assign cells based on the genetic variation between individuals. The count matrices from the Cell Ranger output were analyzed in an R-based Seurat workflow. The count matrix from the published dataset GSE143356 (Guo et al., 2021) was analyzed together with our in-house fetal testis dataset. The fetal male mesonephros/epididymis dataset and fetal ovary dataset were analyzed separately using the same workflow. Briefly, for quality control, cells expressing less than 750 or more than 7000 genes or having more than 50000 UMIs or with more than 10% of the total UMIs coming from mitochondrial genes were excluded from further analysis. Additionally, we also excluded cells with more than 6% of UMIs mapping to dissociation-induced genes (van den Brink et al., 2017). Data was then normalized using the NormalizeData function with the parameter scale.factor set to 50000. To focus on cell type-specific differences in the datasets, batch effects between the datasets were corrected using the fastMNN function from R package batchelor (v1.6.0). The top 2000 (default) highly variable features (genes) were selected with FindVariableFeatures function followed by Principal Component Analysis (PCA). The first 15 principal components (PCs) were used to calculate cell clusters for all datasets, while the resolution parameter was set to 0.5 for the male testis dataset, 0.4 for the male mesonephros/epididymis dataset and 0.2 for the female ovary dataset. Subsequently, cells were projected on a two-dimensional plot using the Uniform Manifold Approximation and Projection (UMAP) algorithm. The R function hclust was used for hierarchical clustering of the fetal testis dataset using all genes. Differentially expressed genes (DEGs) for each cluster were calculated with function FindAllMarkers (only.pos = TRUE) and filtered with pct.1 > 0.6 and p_val_adj < 0.05.

To perform sub-clustering, we first isolated the specific cluster(s) of interest using the subset function from Seurat, followed by the same workflow as above. For rete testis sub-clustering, 20 PCs with resolution 0.5 were applied after FastMNN batch correction (k=5). For rete ovarii sub-clustering, 11 PCs with resolution 0.4 were applied. The 50 most variably expressed genes were selected based on group means using R function rowVars from the package genefilter (v1.74.0) and function heatmap.2 from the R package gplots was used to plot the heatmap.

For differential expression analysis and GO term enrichment, DEGs between groups were first calculated using the function Findmarkers from Seurat, followed by visualization of the results using the EnhancedVolcano (v1.10.0) function. After filtering for p_val_adj <0.05 and avg_log2FC >0.5, DEGs were annotated using the AnnotationDbi (v1.54.1) package and imported for GO term enrichment analysis using the enrichGO function from the DOSE (v3.18.1) package. The GO terms were called specifically for biological process and filtered for p_val_adj <0.05.

### Immunofluorescence

Human material was fixed in 4% PFA overnight at 4°C in a rocking platform, washed in PBS and stored in 70% ethanol at 4°C. The tissue was embedded in paraffin, using a Shandon Excelsior Tissue processor (Thermo Scientific). With a RM2065 Microtome (Leica Instruments), paraffin blocks were sectioned and the sections (5μm thick) were transferred onto StarFrost slides (Knittel) and rested overnight at 37°C. Thereafter, the sections were deparaffinized in xylene (2x 10 min) and rehydrated in decreasing ethanol concentration steps (2x 100%, 90%, 80%, 70%) and finally washed in distilled water. Antigen retrieval was performed in a microwave for 20min at 98°C using 0.01M sodium citrate buffer (pH 6.0) for all primary antibodies, except for GATM, PAX8 and FOXL2 primary antibodies that required antigen retrieval in 10mM Tris/1mM EDTA buffer (pH 9.0). Afterwards, the sections were cooled down to room temperature (RT), washed 2x in PBS for 5min, 1x in PBST (0.05% Tween-20 in PBS) for 5min and blocked using blocking buffer (1% BSA, 0.05% Tween-20 in PBS) in a humidified chamber for 1 hour at RT. Next, the sections were incubated with primary antibodies diluted in blocking buffer overnight at 4°C in a humidified chamber, washed twice in PBS and once in PBST (5min each wash) and incubated with secondary antibodies and DAPI diluted in blocking buffer, in a humidified chamber for 1 hour at RT. Sections were then washed twice in PBS, once in PBST and once in distilled water (5min each wash) at RT and mounted using ProLong Gold. All primary and secondary antibodies as well as the working dilutions used are listed in the Key Resource Table.

### Image analysis

Sections used for immunofluorescence were imaged on a Leica TCS SP8 confocal microscope (Leica) with LAS X software (Leica) using the HC PL APO 40x/1.40 oil objective. Overview scan of all slides were taken with ZEISS Axioscan 7 slide scanner (ZEISS Group). Acquired images were further analyzed with ImageJ software.

## SUPPLEMENTAL TABLES

**Table S1. Differentially expressed genes in the different cell clusters of human fetal testis**

**Table S2. Differentially expressed genes in the different cell clusters of human fetal male mesonephros and epididymis**

**Table S3. Common and specific marker genes in fetal rete testis epithelial cells and feal male mesonephros/epididymis epithelial cells**

**Table S4. Enriched GO terms for common and specific genes in fetal rete testis epithelial cells and feal male mesonephros/epididymis epithelial cells**

**Table S5. Differentially expressed genes in the different cell clusters human fetal ovary**

