## Supplementary material for "Characterization of the human fetal rete region by single cell transcriptional analysis of gonads and mesonephros/epididymis": Suppl Figure S1-S3

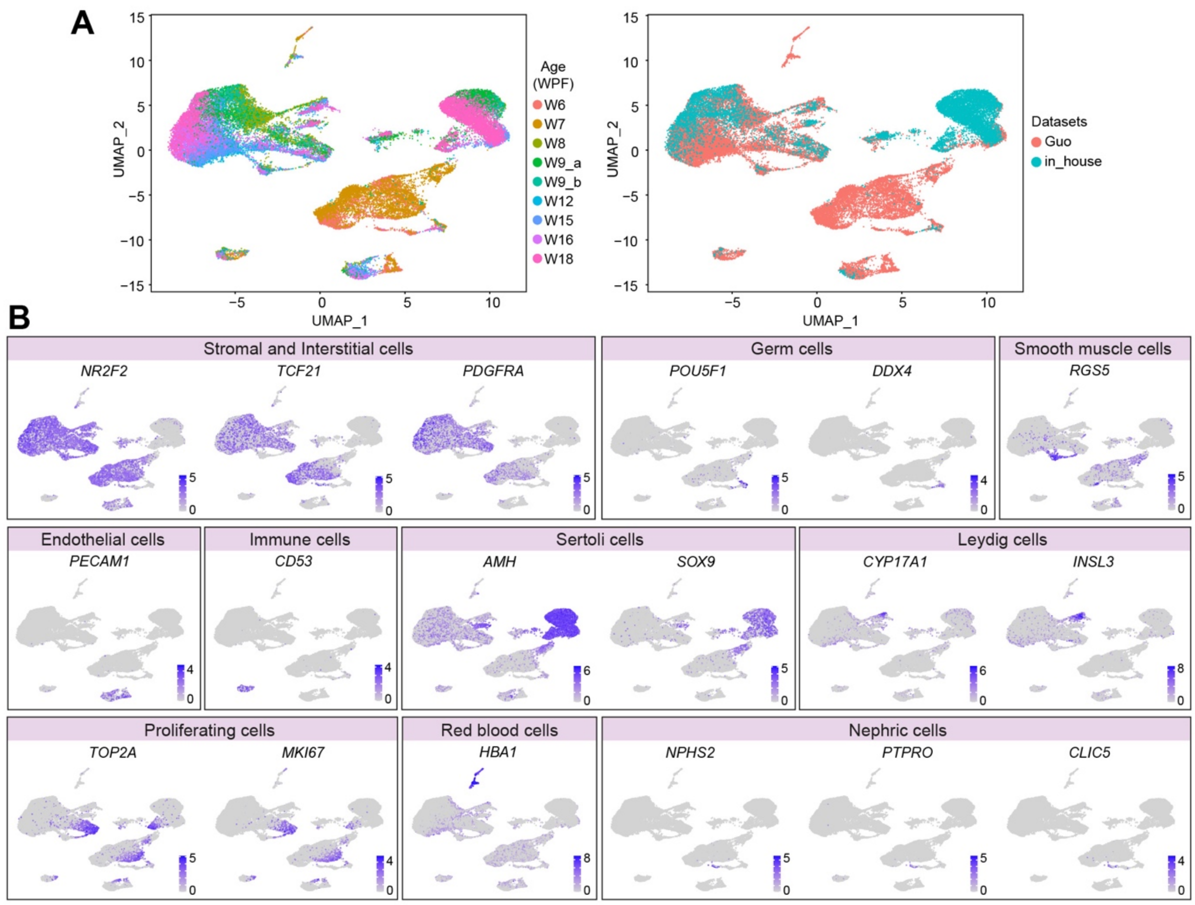


**Figure S1. Identification of cell clusters of human fetal testis**

(**A**) Uniform manifold approximation and projection (UMAP) plot showing cell clusters obtained for the human fetal testis depicting the developmental age (WPF) (left panel) and the origin of the datasets (right panel). (**B**) UMAP plot showing expression of selected markers for the major cell types in the fetal testis.

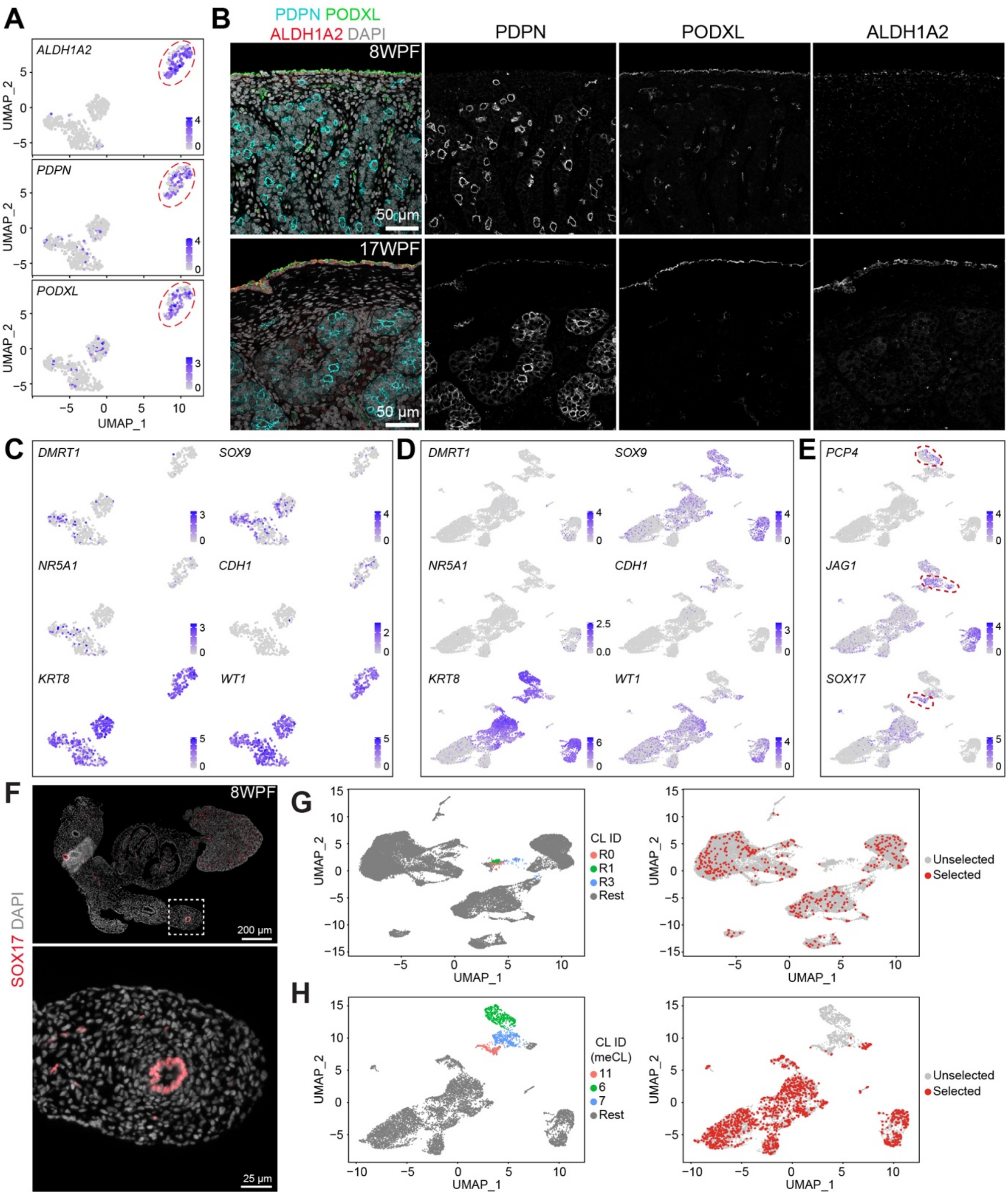


**Figure S2. Characteristics and selection of specific fetal cell populations of interest**
(**A**) Uniform manifold approximation and projection (UMAP) plot from the sub-clustering of mCL7 depicting expression of *ALDH1A2*, *PDPN* and *PODXL*, which mark surface epithelial cells (clusters R2 and R4). Red dashed circles indicate cell clusters R2 and R4. (**B**) Immunofluorescence for ALDH1A2, PDPN and PODXL in 8WPF and 17WPF fetal testes. Scale bars are 50 µm. (**C**) UMAP plot showing sub-clustering of mCL7 depicting expression of selected markers. (**D**) UMAP plot from the human fetal mesonephros and epididymis depicting expression the selected markers shown in C. (**E**) UMAP plot from the human fetal mesonephros and epididymis depicting expression of *PCP4*, *JAG1* and *SOX17*. **(F)** Immunofluorescence for SOX17 in 8WPF female ovary and mesonephros. White dashed box

in the overview image indicates the magnified region bellow. Scale bars are 200 µm in the overview and 25 µm in the magnification. (**G**) UMAP plot showing the rete epithelial cells (R0, R1, R3) and the rest of the cells from the dataset (left panel) and the randomly selected cells from the rest of the dataset for differentially expression analysis (right panel). (**H**) UMAP plot showing the mesonephric epithelial cells (meCL6, meCL7, meCL11) and the rest of the cells from the dataset (left panel) and the randomly selected cells from the rest of the dataset for differentially expression analysis (right panel).

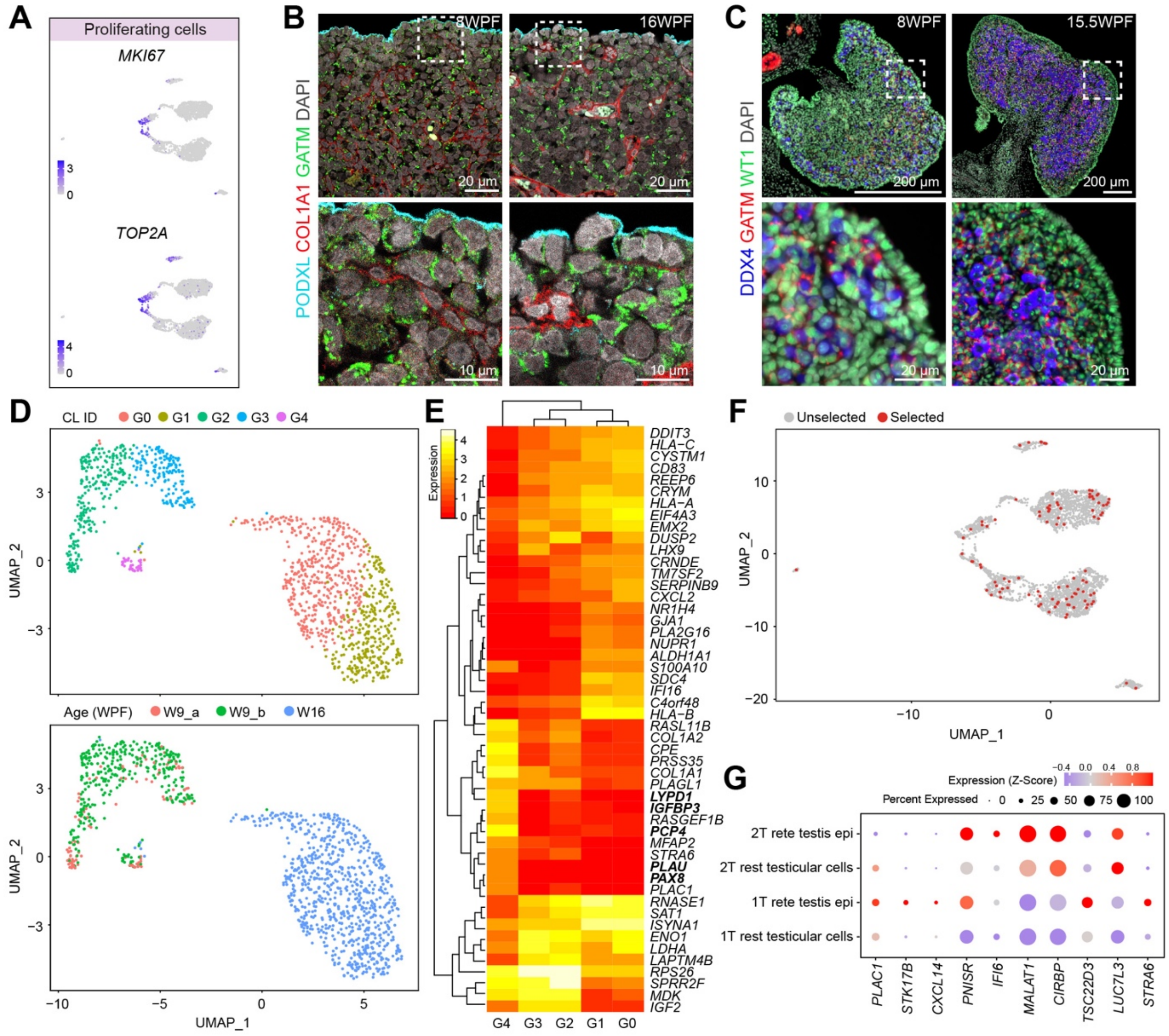


**Figure S3. Identification of specific cell populations in fetal ovaries**
**(A)** Uniform manifold approximation and projection (UMAP) plot from the fetal ovarian cells depicting expression of *MKI67* and *TOP2A*. **(B)** Immunofluorescence for PODXL, COL1A1 and GATM in 8WPF and 16WPF fetal ovaries. White dashed boxes in the overview image (top panel) indicate the magnified region in the bottom panel. Scale bars are 20 µm in overview and 10 µm in the magnification. **(C)** Immunofluorescence for GATM, WT1 and DDX4 in 8WPF and 16WPF fetal ovaries. White dashed boxes in the overview image (top panel) indicate the magnified region in the bottom panel. Scale bars are 200 µm in overview and 20 µm in the magnification. **(D)** UMAP plot showing the sub-clustering of fCL0 (top panel) and developmental age/donor (bottom panel). **(E)** Heatmap showing the top 50 most variably expressed genes in the sub-clusters of fCL0 (G0, G1, G2, G3, G4). **(F)** UMAP plot showing selected cells from the rest of the dataset other than rete ovarii cells for differential expression analysis. **(G)** Dot plots showing the scaled expression (Z-score) of the 10 genes specifically upregulated in the rete ovarii epithelial cells in first and second trimester rete testis epithelial cells and the rest of the cells in the testis.
